# Asymmetric Activation of Retinal ON and OFF Pathways by AOSLO Raster-Scanned Visual Stimuli

**DOI:** 10.1101/2024.12.17.628952

**Authors:** Sara S. Patterson, Yongyi Cai, Qiang Yang, William H. Merigan, David R. Williams

**Affiliations:** Flaum Eye Institute, University of Rochester Medical Center, Rochester, NY, 14642; Del Monte Institute for Neuroscience, University of Rochester Medical Center, NY, 14642; Institute of Optics, University of Rochester, Rochester, NY, 14627; Center for Visual Science, University of Rochester, Rochester, NY, 14627

## Abstract

Adaptive optics scanning light ophthalmoscopy (AOSLO) enables high-resolution retinal imaging, eye tracking, and stimulus delivery in the living eye. AOSLO-mediated visual stimuli are created by temporally modulating the excitation light as it scans across the retina. As a result, each location within the field of view receives a brief flash of light during each scanner cycle (every 33-40 ms). Here we used *in vivo* calcium imaging with AOSLO to investigate the impact of this intermittent stimulation on the retinal ON and OFF pathways. Raster-scanned backgrounds exaggerated existing ON-OFF pathway asymmetries leading to high baseline activity in ON cells and increased response rectification in OFF cells.

## 1. Introduction

Adaptive optics (AO) ophthalmoscopy provides diffraction-limited resolution by detecting and correcting for the eye’s monochromatic aberrations in real time (Liang et al., 1997; Williams et al., 2023). While adaptive optics can be combined with many forms of ophthalmoscope; the combination of AO and confocal scanning laser ophthalmoscopy (AOSLO) has become a particularly valuable for vision research (Webb and Hughes, 1981; Roorda et al., 2002; Roorda, 2011). Rapid online eye-tracking for AOSLO can be used to make highly accurate measurements of eye movements and for online stabilization of visual stimuli on the cone mosaic (Arathorn et al., 2007; Ratnam et al., 2017). The precision of AOSLO-mediated stimulus delivery has enabled unprecedented experiments exploring structure-function relationships in both health and disease (Sincich et al., 2009; Tuten et al., 2012; Sabesan et al., 2016; McGregor et al., 2020).

Stimulus delivery through an AOSLO differs from most modern stimulators used in visual neuroscience in that each “frame” is not presented instantaneously. Instead, spatial patterns are “written” directly into the raster scan by temporally modulating the laser’s intensity as it sweeps across the field of view (FOV) (Poonja et al., 2005). The frame rate, typically 25-30 Hz, is defined by the time taken for the slow scanner to move the fast scan line across the full FOV. In this respect, AOSLO stimulation resembles raster-scanned cathode-ray tube (CRT) displays (Bach et al., 1997). However, what distinguishes AOSLO raster-scanning is the precise focusing of the excitation light provided by adaptive optics and the absence of the phosphor persistence characteristic of CRT displays. A consequence of this precision is that each location within the FOV receives a single brief pulse of focused light within each frame. The physiological impact of this specific form of periodic intermittent stimulation is unclear; however, psychophysical studies have demonstrated that visual perception can change in unexpected ways when periodic stimuli have extreme light-dark ratios (Cobb, 1934; McNemar, 1951). Some of these effects may arise within the circuitry of the retina.

Here we asked whether and how the temporal pattern of excitation created by AOSLO-mediated stimulus delivery impacts the physiology of macaque foveal retinal ganglion cells (RGCs). We utilized fluorescence AOSLO to simultaneously deliver visual stimuli to the foveal cones while imaging the responses of downstream RGCs expressing the calcium indicator GCaMP6s (Yin et al., 2014; Godat et al., 2022; McGregor et al., 2018) (**Figure 1B**). We compared the responses of RGCs to stimuli delivered through the AOSLO with a 561 nm laser to those delivered by a Maxwellian view stimulator independent from the AOSLO (**Figure 1A**). Non-invasive retinal physiology with AO and calcium imaging is a relatively new technique and the extent to which the responses obtained are comparable to more common retinal physiology techniques has not been systematically investigated. While quantitative differences are inevitable given the many technical details that differ between *in vivo* calcium imaging and *ex vivo* electrophysiology, we would expect the results to remain qualitatively comparable. By exploring the impact of AOSLO-mediated stimulus delivery, we sought to test this assumption.

**Figure 1.**
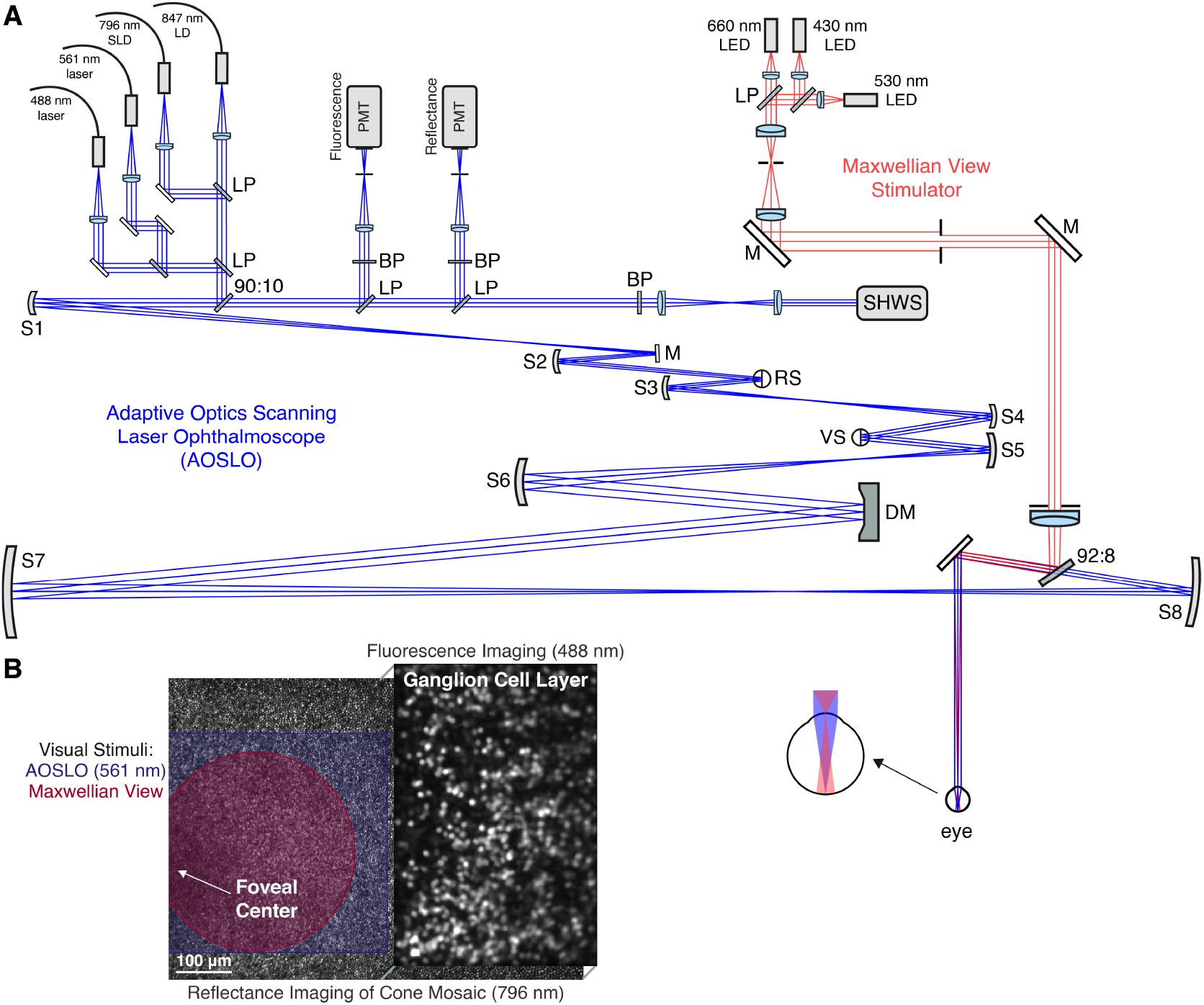
**(A)** System diagram of the AOSLO (blue) and the Maxwellian view (red). The AOSLO contains three key pupil conjugates at the deformable mirror (DM), the resonant scanner (HS) and the galvanometric scanner (VS). Additional components include spherical mirrors (S1-S8), flat mirrors (M); long-pass filters (LP), band-pass filters (BP), Shack-Hartmann wavefront sensor (SHWS) and photomultiplier tubes (PMT). **(B)** Diagram illustrating the spatial arrangement of each visible light source. Reflectance imaging of the cone mosaic at 796 nm was performed at across the full 3.69 x 2.70° field of view while a 488 nm laser was focused on the ganglion cell layer and restricted to either the left or right side of the field of view. Visual stimuli were presented to the foveal cones on the opposite side, either through the AOSLO with a 561 nm laser or independent from the AOSLO with the 3-LED Maxwellian view stimulator. All experiments were performed in the fovea where RGCs are displaced laterally from their cone inputs.

**Figure 2.**
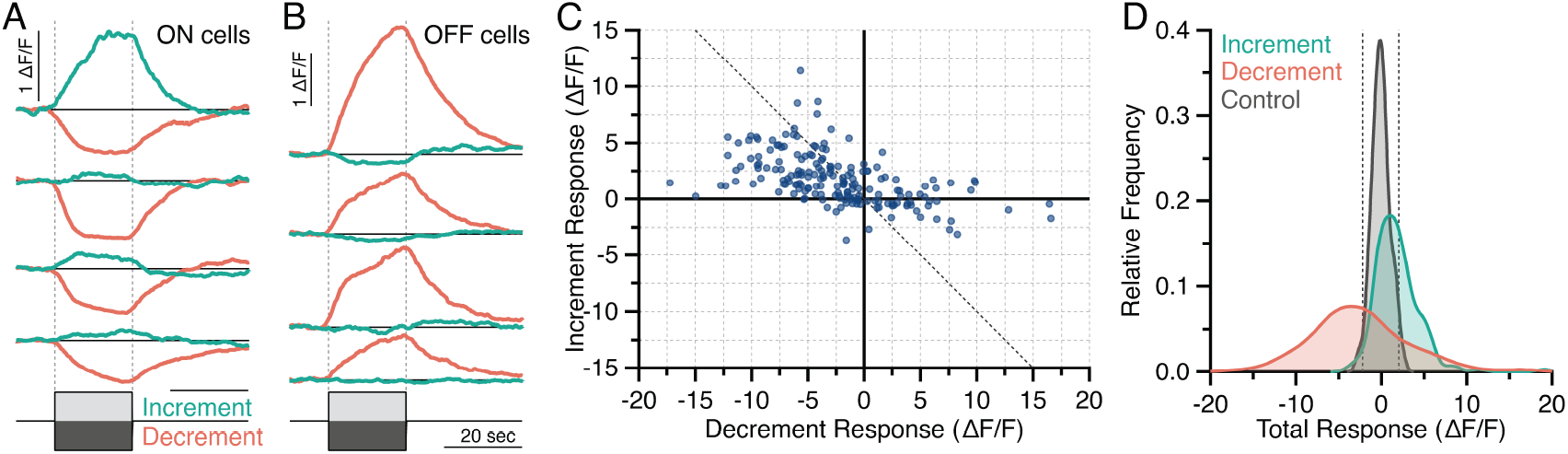
**(A-B)** Representative responses to 20-second contrast increments (green) and decrements (red) for 4 ON cells (A) and 4 OFF cells (B). The gray vertical dashed lines indicate the start of the contrast increment/decrement and the black horizontal lines mark the baseline (0% ΔF/F) for each cell. All vertical scale bars are 50% ΔF/F and all horizontal scale bars are 20 seconds. **(C)** Comparison of the total stimulus responses to the contrast increment and decrement for 187 cells. The gray dashed line indicates equal and opposite increment and decrement responses. **(D)** Kernel density estimates summarizing the distribution of total stimulus responses to the contrast increment, decrement, and a control stimulus (0% contrast). The vertical dashed lines indicate the response significance cutoff which was set at 2 SDs from the mean total response for the control stimulus.

Our focus in comparing calcium responses with *ex vivo* electrophysiology data was on the responses of ON and OFF RGCs to increment and decrement stimuli. These simple stimuli are among the first used when classifying RGCs because they reveal fundamental information about response polarity – that is, whether an RGC has ON, OFF or ON-OFF responses (Kuffler, 1953; Ichinose and Habib, 2022). How these responses arise from and relate to the underlying retinal circuitry is well understood, making them a strong benchmark for evaluating RGC response properties under non-standard stimulation and measurement conditions. For example, the balance between the ON and OFF pathways is thought to impact the efficacy of bipolar cell stimulation by retinal prosthetics (Carleton and Oesch, 2024) and may be altered in conditions like myopia and amblyopia (Pons et al., 2019; Poudel et al., 2024).

Before proceeding, we will briefly summarize the expected response properties of ON and OFF RGCs to contrast increments and decrements relative to a photopic background light level. All RGCs have a limited dynamic range bounded by the rectification of responses below spike threshold and the saturation of responses near the maximal spike rate. The visual system optimizes contrast sensitivity by dedicating the limited dynamic range of ON and OFF RGCs to signaling increments and decrements, respectively (Gjorgjieva et al., 2014; Sterling and Laughlin, 2017). Although often thought of as differing only in the sign of their response, ON and OFF RGCs do not have mirror symmetric responses to increments and decrements (**Figure 3A**). In cone-mediated photopic vision, OFF cells respond only to decrements, whereas ON cells respond to both increments and low-contrast decrements (Chichilnisky and Kalmar, 2002; Zaghloul et al., 2003). This ON-OFF asymmetry is rooted in the different baseline spike rates of ON and OFF RGCs, which are largely inherited from presynaptic ON and OFF bipolar cells (Zaghloul et al., 2005; Margolis and Detwiler, 2007; Liang and Freed, 2010). Most OFF RGCs are more hyperpolarized than their ON counterparts and exhibit very low spontaneous discharge (near 0 spikes/sec at photopic light levels), rendering them unable to signal increments with further reductions in spike rate. In other words, the output of the retinal OFF pathway is strongly rectified. In contrast, ON RGCs have a higher baseline spike rates and can signal decrements through decreases in spiking, though the majority of their dynamic range is devoted to increments. While some ON-OFF RGC pairs break this pattern (Ravi et al., 2018), these ON-OFF asymmetries are largely conserved across mammalian RGCs, including ON and OFF parasol and midget RGCs, the most common primate RGC types (Chichilnisky and Kalmar, 2002; Soto et al., 2020; Percival et al., 2022). This study investigates whether and how these characteristic ON and OFF RGC responses are altered by *in vivo* calcium imaging and AOSLO-mediated stimulus delivery.

## 2. Methods

### 2.1. Animal care

Two macaques (*Macaca fascicularis*), two female and one male, were housed in pairs in an AAALAC accredited facility. All animals were in the care of the Department of Comparative Medicine veterinary staff of the University of Rochester’s Medical Center, including several full-time veterinarians, veterinary technicians, and an animal care staff who monitored animal health. Additional details on the vivarium environment are detailed in our previous work (McGregor et al., 2020; Godat et al., 2022). This study was carried out in strict accordance with the Association for Research in Vision and Ophthalmology (ARVO) Statement for the Use of Animals and the recommendations in the Guide for the Care and Use of Laboratory Animals from the National Institutes of Health. The animal protocol was approved by the University Committee on Animal Resources (UCAR) of the University of Rochester under PHS assurance number D16-00188 (A3292-01). Each macaque had no known history of ocular disease or vision abnormality.

### 2.2. Viral delivery

Intravitreal injections of *AAV2:CAG:GCaMP6s* were carried out in each macaque, as previously described (Yin et al., 2011; Chen et al., 2013). Briefly, each eye was sterilized with 50% diluted Betadine, and the injection was made in the middle of the vitreous approximately 4 mm behind the limbus using a tuberculin syringe and 30-gauge needle. Following injection, each eye was imaged with a conventional scanning light ophthalmoscope (Heidelberg Spectralis) using the 488 nm autofluorescence modality to determine onset of GCaMP expression in the ganglion cell layer (GCL) and to monitor eye health. Both macaques received subcutaneous injections of Cyclosporine A prior to and after injection of viral vectors to suppress the immune response and promote strong, maintained expression. The data in this study was collected 1.7 years post-injection from the right eye of M3 and between 0.8-2 years post-injection from both eyes of M4. Additional information regarding the immune suppression and injection history for M3 and M4 can be found in our prior work (Godat et al., 2024).

GCaMP6s expression under the promoter CAG labeled foveal neurons throughout the ganglion cell layer, 1-2% of which are estimated to be displaced amacrine cells (Curcio and Allen, 1990). By focusing on large, population-level trends, the results of the present study are best interpreted as representing the responses of foveal RGCs, the majority (∼86%) of which are ON or OFF midget RGCs (Peng et al., 2019).

### 2.3. Imaging preparation

Anesthesia and animal preparation procedures have been previously published (McGregor et al., 2020) and are briefly summarized here. The anesthesia and preparation were performed by a veterinary technician licensed by the State of New York (USA). All animals were fasted overnight prior to anesthesia induction the morning of an imaging session. During a session, animals were placed prone onto a custom stereotaxic cart and covered with a Bair Hugger warming system to maintain body temperature. Monitoring devices including rectal temperature probe, blood pressure cuff, electrocardiogram leads, capnograph, and a pulse oximeter, were used to track and record vital signs. Temperature, heart rate and rhythm, respirations and end tidal CO_2_, blood pressure, SPO2, and reflexes were monitored and recorded every fifteen minutes. Pupil dilation was accomplished using a combination of 1% Tropicamide and 2.5% Phenylephrine. Each eye was fitted with a hard contact lens (AVT) to relieve adaptive optics from the need to correct lower order aberrations and to keep the eye from drying out during the imaging session.

### 2.4. AOSLO retinal imaging

Data was collected using a fluorescence AOSLO, described previously (Gray et al., 2006; Hunter et al., 2011; Godat et al., 2022) and diagrammed in **Figure 1A**. An 847 nm diode laser source (QPhotonics) was used as a beacon for the Shack-Hartmann Wavefront Sensor to measure optical aberrations in each animal’s eye in real time at ∼20 Hz. A deformable mirror (AL-PAO) with 97 actuators was used to correct those aberrations in a closed loop at approximately 8 Hz across a 7.2 mm diameter pupil.

A 796 nm superluminescent diode (Superlum) was focused on the foveal cone photoreceptor mosaic to collect images (using a ∼ 2 Airy disk confocal image, 20 µm) used for navigating to the same retinal location in each experiment and for offline motion correction/registration of fluorescence images. A 488 nm laser (Qioptiq) was focused on the GCL and used to excite fluorescence from GCaMP6-expressing cells, which was detected through a 520/35 nm filter (using a ∼2 Airy disk confocal pinhole, 20 µm in M3, 25 µm in M4). The 488 nm excitation laser was presented only during forward raster scans of the AOSLO and only to the portion of the imaging field where RGCs were present (i.e., laterally displaced from the foveal cones and axially displaced to focus on the GCL; **Figure 1B**). The somas of the foveal RGCs imaged lay at the margins of the foveal pit, approximately 1-4° from the foveal center. RGCs were given at least 5 minutes to adapt to the presence of the 488 nm laser before data collection began. During this time, the X and Y position of the fluorescence PMT was empirically adjusted to maximize signal strength. This procedure fine-tuned the source and PMT position optimization performed for the 796 nm, 561 nm and 488 nm sources prior to each experiment using a model eye (Godat et al., 2022).

All data were collected with a 3.69 x 2.70° FOV, with the exception of the contrast response functions in **Figure 3B-C** which were collected at a 3.37 x 2.72°FOV. To obtain accurate estimates of the FOV, the AOSLO system was calibrated using an 80 lp/mm Ronchi ruling (R1L3S15N, ThorLabs) to obtain a lookup table for translating scanner control voltages into FOV measurements in degrees of visual angle.

### 2.5. Light safety

The total power at the cornea was measured before each experiment to ensure light safety calculations were accurate. All light sources were delivered through a dilated pupil of 6.7 mm. The 488 nm laser was 1.7 µW/cm^2^ and posed the primary limitation on light safety at the large FOVs used in the present study. The 796 nm reflectance imaging source and 847 nm wavefront sensing beacon were 250 µW/cm^2^ and 30 µW/cm^2^, respectively. The reflectance imaging source and the inevitable light leaked from the 561 nm laser provided a constant low photopic background to all experiments. Room lights were turned off during experiments to minimize ambient light.

The total exposure during each experiment was kept below the maximum permissible exposure for human retina according to the 2014 American National Standards Institute and most exposures were also below a reduced limit that was further scaled by the squared ratio of the numerical aperture of the human eye to the primate eye (∼0.78-0.85). In this study, exposures during 2-4 hours of imaging ranged from 60-94% of the human retinal limit, with the majority between 60-85%. All sessions were spaced a minimum of 5 days apart per animal, so cumulative light exposure was not calculated.

### 2.6. AOSLO visual stimuli

AO-corrected, stabilized visual stimuli were presented by a 561 nm laser (iChrome MLE, Toptica) focused on the photoreceptor mosaic. An acousto-optic modulator (AOM; Brimrose Corporation) was used to create patterned stimuli by modulating the laser’s output temporally as it scanned across the FOV (Poonja et al., 2005). The frame rate was ∼ 25.3 Hz and, at the FOVs used in the present study each pixel was approximately 1.5 µm^2^. Stimuli were specified as 256×256 8-bit videos where each pixel value determined the AOM modulation of the 561 nm laser. Stimuli were stabilized on the cone mosaic using a reference reflectance image of the cone mosaic taken at the beginning of each experiment (Yang et al., 2014).

Both the AOSLO stimuli and the Maxwellian view stimuli described below were presented following a 20 second baseline period and at least five minutes of adaptation time was provided any time the mean light level changed before obtaining responses to increment/decrement stimuli. Increments and decrements relative to a background light level were described by percent Weber contrast: 100 * (*I*_*stim*_ − *I*_*mean*_)*/I*_*mean*_ where *I*_*mean*_ is the intensity of the background and *I*_*stim*_ is the intensity of the increment or decrement.

### 2.7. Maxwellian view visual stimuli

A Maxwellian view stimulator with three LEDs (M430L3, M530L3 and M660L4, ThorLabs) presented a spatially-uniform 1.25 degree diameter, spatially-uniform circle to the foveal cones (Leibowitz, 1954; Baron, 1973). The light paths from the Maxwellian view and the AOSLO were combined after the final spherical mirror using a 92:8 pellicle (BP208, ThorLabs). The three LEDs’ voltages were updated at 500 Hz (LEDD1B, ThorLabs). The voltages of each LED were recorded every 2 ms and also at the beginning of each AOSLO frame for synchronization with the calcium imaging data. The LEDs were calibrated using a spectrophotometer (STS-VIS, Ocean Optics) validated by a NIST traceable blackbody source as detailed in our previous work (Godat et al., 2022, 2024). Calibrations were performed after each system alignment and/or whenever the pellicle beamsplitter’s position was adjusted.

While the two stimulators produced comparable power at the cornea, we did not equate their efficacy at driving the L-, M- and S-cones, which nonetheless preserved the manner in which these two methods for light delivery are typically used. For 3-primary light sources, contrast modulations are typically performed around equal energy white to avoid placing the L-, M- or S-cones in an uncommon adaptation state. Both stimulators were ∼ 1.2x more effective for L-cones than M-cones, but the blue LED primary produced greater activity in S-cones. Given the rarity of S-cones and RGCs with strong S-cone input in the macaque fovea (Godat et al., 2024), a difference in S-cone activation is unlikely to account for the large, population-level differences between the two stimulators.

Unlike the spatial stimuli presented through the AOSLO, the Maxwellian view stimulus was not stabilized on the retina. While some residual eye motion is present even in anesthetized, paralyzed macaques due to head motion during respiration, the light falling on the RGC receptive fields remains spatially homogeneous when the receptive fields are well-centered within the 1.25-degree stimulus region. At the beginning of each experiment, we confirmed that the Maxwellian view stimulus was centered on the RGCs’ receptive fields by checking for large changes in fluorescence in phase with eye motion due to respiration, which were caused by the RGCs’ receptive field moving across the edge of the Maxwellian view stimulus.

### 2.8. Data analysis

Fluorescence videos were co-registered with the simultaneously-acquired reflectance videos frame-by-frame with a strip-based cross-correlation algorithm (Yang et al., 2014). Registration was performed relative to a reference image of the cone mosaic, taken during the experiment and used for online stabilization. Because small changes in pupil position during the course of an experiment could lead to small offsets in registered videos, a secondary registration using SIFT (Lowe, 2004) was performed on the summed Z-projections of each fluorescence video. The summed Z-projections were also used for manual segmentation of regions of interest (ROIs) using ImageJ’s ROI Manager. All subsequent analyses were performed in MATLAB 2024a (Mathworks, RRID: SCR_001622) and Igor Pro 9 (Wavemetrics, RRID: SCR_000325). Final figures were created in Igor Pro or Adobe Illustrator (Adobe, RRID: SCR_010279).

Calcium responses were extracted as the average fluorescence of all pixels within an ROI. Fluorescence responses (*F* ) were baseline-corrected by calculating the ΔF/F as follows:

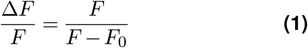

where *F*_0_ was the average fluorescence during the 10 seconds preceding stimulus onset during which the mean background light level was present. The ΔF/F traces were smoothed with a 4-second moving window in all analyses except the temporal correlation coefficient analysis in **Figure 9** where a four-fold lower window was used to preserve temporal resolution. Responses were normalized for **Figure 3B** and the correlation coefficient analyses in **Figure 9** by dividing the full ΔF/F response by the peak response metric described below. Cells from different experiments were only combined for analyses where responses were normalized. The total response metric took the approximate integral (MATLAB’s trapz function) of the response within the stimulus window divided by the stimulus time in seconds. The peak response metric used in **Figures 7B** and **7E** reflects the 2nd or 98th percentile of the ΔF/F response, depending on which had a greater absolute value. Percentiles were used instead of minimum and maximum values to limit the impact of noise. For responses that were frequently biphasic (e.g., the contrast decrement-increment stimulus in **Figure 5E** and the intensity increments in **Figure 8B**), we calculated the amplitude (98th percentile minus the 2nd percentile) rather than the peak response. The rectification index was computed as the 98th percentile divided by the amplitude.

**Figure 3.**
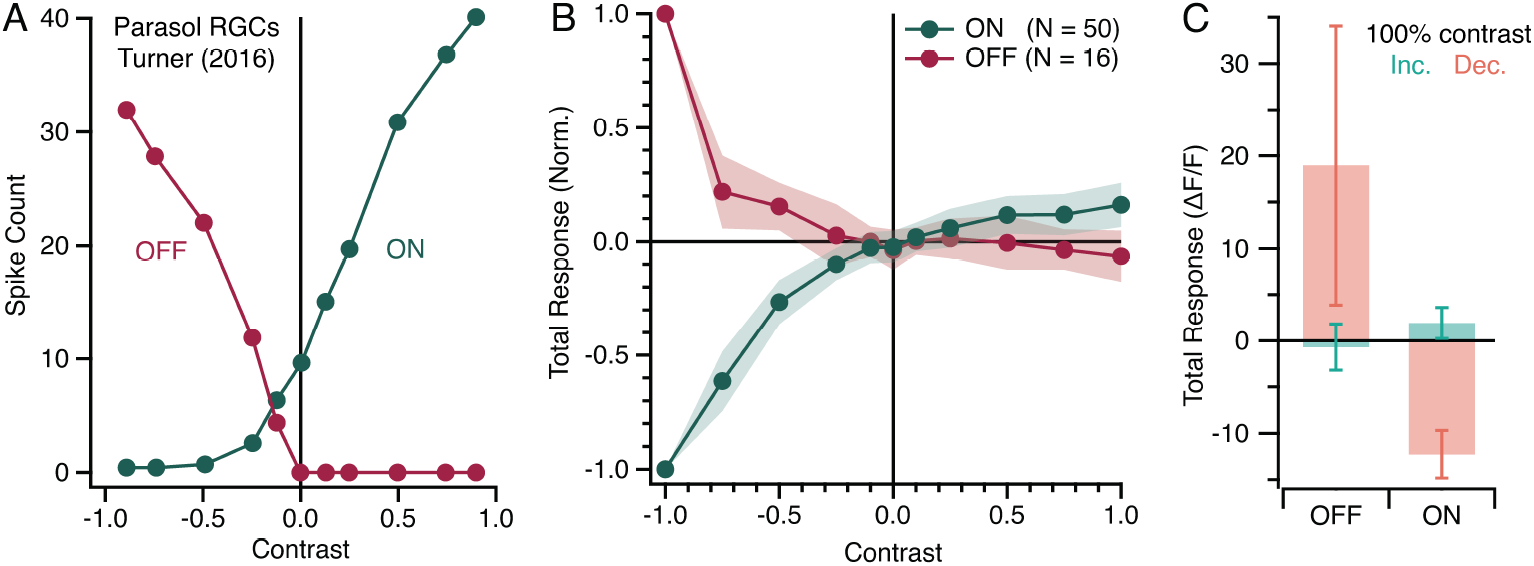
**(A)** Representative contrast response functions (CRFs) obtained with ex vivo electrophysiology from ON (green) and OFF (red) parasol RGCs (Turner and Rieke, 2016). **(B)** Average CRFs for ON and OFF measured at 0%, 10%, 25%, 50%, 75% and 100% contrast. The mean and shaded standard deviation were calculated from the total response during each 20 second contrast step for 16 OFF cells and 50 ON cells. Each cell’s CRF was individually normalized before averaging. **(C)** Mean and standard deviation of the absolute total stimulus responses for 100% contrast increments and decrements for 16 OFF and 50 ON cells.

All summary statistics are listed as mean ± SD, unless otherwise specified. The Hartigan’s dip test was used to test for bimodality and the two-sample Kolmogorov-Smirnov test was used to compare distributions from ON and OFF cells. All other statistical tests were two-sample t-tests unless otherwise indicated. For visual clarity, overlapping distributions were plotted as empirical cumulative distribution functions or kernel density estimates using bandwidths determined by the normal-approximation method.

## 3. Results

### 3.1. Unexpected responses of ON and OFF cells to increments and decrements delivered via AOSLO raster scanning

We began by characterizing the light responses of GCaMP6s-expressing neurons in the foveal ganglion cell layer to spatially uniform contrast increments and decrements relative to a photopic mean light level (referred to as the “background light”). The background light was present before and after each increment/ decrement and retinal neurons were given at least five minutes to fully adapt to any change in background light level. Representative responses from 8 neurons (4 ON and 4 OFF) to 100% Weber contrast increments and decrements are shown in **Figure 2A-B**. As expected, positive responses (increases in ΔF/F) were observed for ON cells during the increment and for OFF cells during the decrement. Unexpectedly, the ON cells also exhibited large negative responses (decreases in ΔF/F) during the decrement stimulus. While ON cells typically have higher baseline spike rates than OFF RGCs that enable some capacity for signaling decrements through decreases in spiking (Margolis and Detwiler, 2007; Percival et al., 2022), the majority of their dynamic range is devoted to increments. The primary decrement response in ON cells should be an increase in ΔF/F at the offset of the decrement, not a decrease in ΔF/F at the onset of the decrement.

To assess these trends over the full population, we quantified the increment and decrement responses by computing a “total stimulus response” metric that took the approximate integral of the ΔF/F values during the increment/decrement divided by the stimulus duration in seconds. **Figures 2C-D** compare the total stimulus response to the increment and decrement for 187 cells. As expected, OFF cells clustered along the positive x-axis in **Figure 2C** because their responses were strongly rectified, with minimal decrease in ΔF/F during the increment. By contrast, ON cells were located in the top left quadrant because they exhibited strong, opposing changes in fluorescence to the increment and decrement. Most ON cells fell below the dashed line in **Figure 2C**, indicating that the decrease in ΔF/F during the decrement exceeded the corresponding increase in ΔF/F during the increment. As above, ON RGCs have some capacity to decrease their activity below baseline to signal decrements; however, the magnitude of the ON cells’ fluorescence decreases in response to the contrast decrement is inconsistent with the response properties of the most common ON primate RGCs.

For reference, we also calculated the total response metric for a control stimulus without increments or decrements (-0.07 ± 1.05 ΔF/F, n = 187; **Figure 2D**). Significant increment/decrement responses were defined as those exceeding two standard deviations above or below the average control stimulus response (dashed lines in **Figure 2D**). 78.87% (140/187) of the decrement responses were significant, but only 41.71% (78/187) of the increment responses. Surprisingly, the ON cells’ negative responses dominated the decrement response distribution; only 37 of the 140 significant decrement responses were positive responses from OFF cells. This is result is unexpected as our stimulus (100% contrast decrements at photopic light levels) should elicit robust responses in retinal OFF cells. While ON and OFF cells are found in different regions of the ganglion cell layer and some ON:OFF sampling bias could be present in our results (Perry and Silveira, 1988), the bias towards ON cells observed here was far more pronounced than in any of our prior studies (McGregor et al., 2018; Godat et al., 2022, 2024). This apparent lack of strong decrement responses in OFF cells will be revisited in later sections.

### 3.2. Contrast response functions of ON and OFF cells

To diagnose the origins of the unexpected responses in **Figure 2**, we next measured contrast response functions (CRFs) across five positive and negative contrast levels (10%, 25%, 50%, 75% and 100%). Stimulus-response functions like CRFs can provide valuable insight into underlying neural activity. For example, the ON-OFF asymmetries in baseline activity and rectification introduced earlier are clear in the CRFs from ON and OFF primate parasol RGCs shown in **Figure 3A** (Turner and Rieke, 2016). For example, ON cells are relatively depolarized at rest and have a non-zero baseline spike rate (response along the vertical line marking the 0% contrast control stimulus in **Figure 3A**), while OFF cells are relatively hyperpolarized and have minimal baseline activity.

The average CRFs for 56 ON cells and 20 OFF cells are shown in **Figure 3B**. The most surprising feature of the measured CRFs was the ON cells’ baseline activity level at 0% contrast. When the stimulus was held constant at the background light level and no increments or decrements were presented, the ON RGCs were, on average, at 86.57% of their maximum response (**Figure 3B**). As a result, the majority of the ON RGCs’ dynamic range was devoted to signaling decrements through decreases in fluorescence. Note that while the total response at 0% contrast is near zero for the average ON CRF in **Figure 3B**, this is enforced by the ΔF/F calculation (**Equation 1**), which normalizes each response to the average fluorescence during a pre-stimulus period where only the background light level is present. As a result, ΔF/F responses can obscure high baseline spike rates.

A consequence of high baseline activity in ON cells would be a reduced capacity to signal increments with further increases in fluorescence. Consistent with this prediction, the ON cells’ responses to 100% contrast increments were substantially smaller than the OFF cells’ responses to 100% contrast decrements (ON: 1.54 ± 1.67, n = 56; OFF: 14.34 ± 12.18, n = 20; **Figure 3C**). In addition, the absolute magnitude of the ON cells’ increment response was, on average, just 15.5% of their response to the 100% contrast decrement. The flattening of the ON cells’ CRF at positive contrasts indicates that their responses were saturated, which compromised discrimination between low and high contrast increments. For example, the difference between the 50% and 75% contrast increments was not statistically significant (p = 0.97). This trend is opposite that expected from prior studies, where the rectification imposed by spike threshold instead limits the ON RGCs’ ability to discriminate between low and high contrast decrements (Chichilnisky and Kalmar, 2002) (**Figure 3A**).

The difference between measured and expected OFF CRFs was less pronounced; however, the average OFF CRF in **Figure 3B** exhibited very little responsivity to low contrast decrements. In some OFF cells, only the 100% contrast decrement resulted in a significant response and, at the population level, responses to the 75% contrast decrement were just 0.22x as large as the responses to the 100% contrast decrement (**Figure 3B**). Taken together, these features were consistent with increased hyperpolarization in the OFF pathway, resulting in greater rectification of the OFF cells’ output. If OFF cells were more hyperpolarized and their outputs even more rectified than usual (e.g., **Figure 3A**), the responses to low contrast decrements would sink below spike threshold, leading to a leftward shift in their CRF as observed in **Figure 3B**. It is possible some OFF cells were sufficiently hyperpolarized that they did not even have detectable responses to the 100% contrast decrement, potentially explaining the low numbers of OFF cells identified in **Figures 2, 3** and **4**.

To summarize the results so far, the ON cells’ negative responses during contrast decrements were much larger than expected from previous studies of ON midget and ON parasol RGCs (Chichilnisky and Kalmar, 2002; Soto et al., 2020; Gogliettino et al., 2023). While the ON cells’ polarity was preserved (ΔF/F increased for contrast increments and decreased for contrast decrements; **Figures 2A-B, 3B-C**), their responses signaled decrements with greater fidelity than increments (**Figure 3B**). More generally, these results point to high baseline activity as a potential underlying mechanism for the ON cells’ unexpectedly large ΔF/F decreases during decrements (**Figure 2B**): if the ON RGCs were already responding strongly to the background light and stopped responding when the background light turned off during a decrement, then a large drop in ΔF/F would be expected. By contrast, the OFF cells exhibited stronger rectification and lower responsivity to all but the highest contrast decrements.

### 3.3. Impact of background light level

Having explored stimulus contrast, we next varied the intensity of the background light level, which can have a strong impact on the responses of retinal neurons and visual perception (Kuffler, 1953; Marcos et al., 2008; Borghuis et al., 2018). Background light level can also increase an existing bias within the visual system toward decrements (Bowen et al., 1992; Angueyra et al., 2022), which generate larger neural responses and lower psychophysical detection/discrimination thresholds compared to increments of equal contrast (Bowen et al., 1989; Yeh et al., 2009). This well-documented increment-decrement asymmetry originates in the cone photoreceptors and could contribute the relatively weak responses to contrast increments in our data (**Figures 2D** and **3C**).

We presented 100% contrast increments and decrements at three background light levels (1.4, 1.8, and 4.7 µW; **Figure 4A**). As the background light level increased, the magnitude of the ON cells’ positive responses to increments decreased, while their negative responses to decrements became more pronounced. This can be seen in the representative ON cell shown in **Figure 4B** and in a population of 56 ON cells in **Figure 4C**. At the highest light level, many ON cells exhibited little or no increase in fluorescence during the increment despite having large negative responses during the decrement. One possible explanation for this result is that the ON cells became increasingly depolarized as the background light level rose. As the ON cells’ baseline activity level approached the upper limit of their dynamic range, their capacity to signal increments with further increases in fluorescence was reduced.

**Figure 4.**
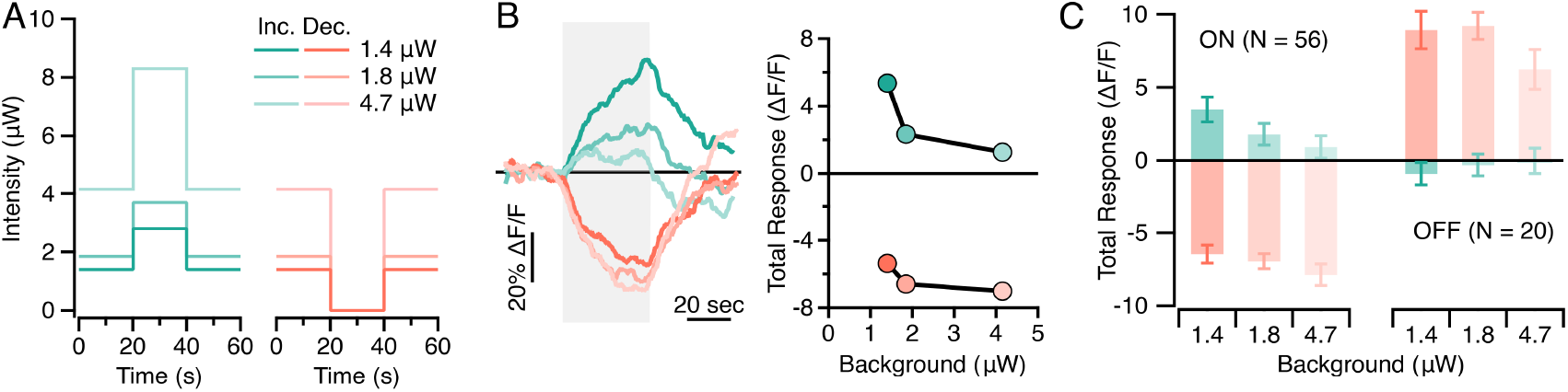
**(A)** Stimulus intensity of 100% contrast increments and decrements at three mean light levels. **(B)** Representative ON cell traces (left) and total stimulus responses (right) to increments and decrements at the three light levels. Colors are the same as used in **A. (C)** The mean and standard deviation of 56 ON and 20 OFF cells’ total stimulus responses to the increment and decrement at each mean light level.

The response profiles of individual OFF cells were more heterogeneous. Some showed increasing or decreasing responses with mean light level while others remained unchanged. At the population level, the magnitude of the OFF cells’ decrement response decreased at the highest light level. Additionally, weak decreases in fluorescence were also observed during the increment, which also declined with increasing light level. This decline in the OFF cells’ negative increment responses indicated the rectification of the OFF pathway increased with mean light level. However, these observations are qualified by the low numbers of OFF cells identified by their positive responses to the decrement stimulus, a trend also seen in the earlier two experiments (**Figure 2-3**).

In summary, the shape of the CRFs in **Figure 3B** indicated an elevated baseline activity in ON cells and increased rectification in OFF cells. The results in **Figure 4C** are consistent with these ideas and indicate a dependence on background light level. Specifically, higher background intensities led to a decreased capacity for signaling increments with ΔF/F increases, consistent with rising baseline activity levels approaching the upper limit of the ON cells’ dynamic range. While there was evidence for greater OFF cell rectification with background light level, these observations are limited by the low numbers of OFF cells identified and the overall heterogeneity of their responses.

### 3.4. Stimulus responses for AOSLO-mediated stimulus delivery vs. a Maxwellian view stimulator

An important implication of the dependence on background intensity observed in **Figure 4** is that the unexpected increment and decrement responses in **Figures 2-3** are stimulus-driven and likely caused by some feature of AOSLO-mediated stimulus delivery. To directly test whether AOSLO-mediated stimulus delivery alters ON and OFF pathway responses, we next compared the ON and OFF cell responses to AOSLO visual stimuli with comparable stimuli presented with a Maxwellian view stimulator (MV). Importantly, the Maxwellian view stimulator is a separate subsystem from the AOSLO and was introduced into the optical path after the final telescope using a pellicle beamsplitter (**Figure 1A**).

To increase the efficiency of data collection in these experiments, we used a decrement-increment stimulus in which the increment immediately followed the decrement (both 20 seconds long and at 100% contrast relative to a photopic background light). **Figures 5A-B** shows representative ON and OFF cell responses to this stimulus when presented through the AOSLO vs. the Maxwellian view. Consistent with our previous findings (**Figures 2A, 3B**, and **4B**), ON cells displayed unexpected decreases in fluorescence during the decrement delivered through the AOSLO. In contrast, the ON cells’ responses to a comparable stimulus presented via the Maxwellian view reflected the rectification expected from *ex vivo* electrophysiology. This difference is evident at the population level in **Figure 5C**, which shows the average normalized responses from 72 ON cells to the AOSLO and Maxwellian view stimuli. Notably, only 3 of the 214 cells analyzed exhibited similar response profiles to both the AOSLO and Maxwellian view stimulus. An example is shown in the bottom trace of **Figure 5A**. These neurons could be displaced amacrine cells that make up 1-2% of the foveal ganglion cell layer (Curcio and Allen, 1990) or rarer RGC types.

**Figure 5.**
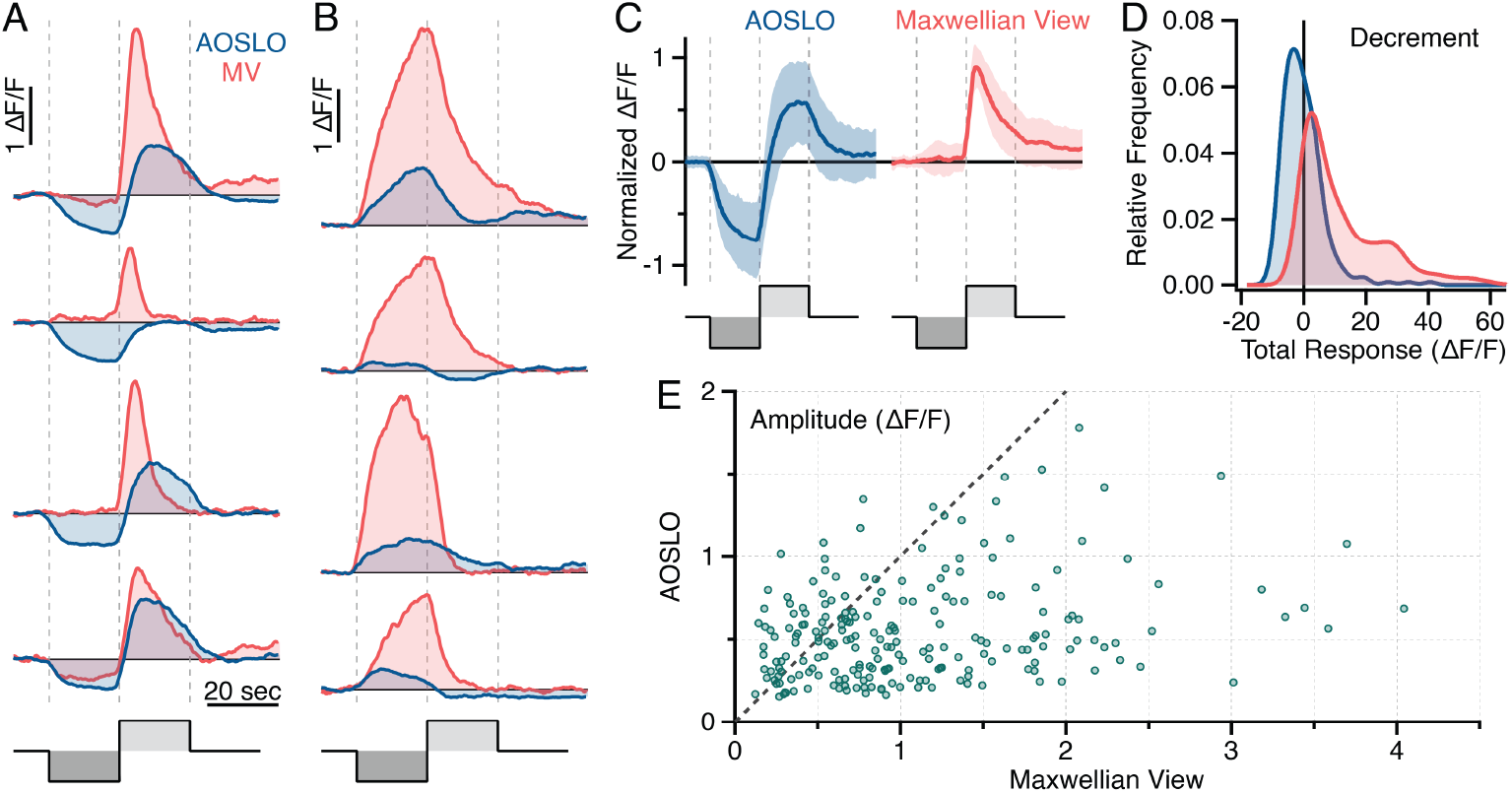
Responses to the decrement-increment stimulus presented through the AOSLO and the Maxwellian view (MV). **(A)** Representative responses from four ON cells to the contrast modulation presented through the AOSLO (blue) and the MV (red). The gray dashed lines mark the onset and offset of the decrement and increment. **(B)** As in A, but showing representative responses from four OFF cells. **(C)** Population-level responses from ON cells to the AOSLO and MV decrement-increment stimulus. The average and standard deviation of the 72 individually normalized traces are shown. **(D)** Kernel density estimates of the total response distribution during the AOSLO and MV contrast decrements (n = 214 cells). **(E)** Comparison of the trough-to-peak amplitudes of responses elicited by the AOSLO vs. Maxwellian view decrement-increment stimulus for the same 214 cells.

Additionally, the total decrement response distributions across the full population of 214 cells in **Figure 5D** revealed a shift towards positive responses in the Maxwellian view condition. The difference in the total decrement responses between the AOSLO and the Maxwellian view was statistically significant (-3.19 ± 6.67 vs. 11.52 ± 13.19, n = 214; p = 2.75e^-39^ ). We confirmed that this result was not dependent on integration over the full 20-second decrement by recalculating the total decrement response over shorter time intervals. Even when measured over the first 3, 5, and 10 seconds of the decrement, the difference between the AOSLO and the Maxwellian view remained statistically significant (10 seconds: -2.08 ± 5.92 vs. 7.56 ± 9.07, p = 7.54e^-33^ . 5 seconds: -1.25 ± 4.62 vs. 4.80 ± 5.86, p = 2.78e^-28^ ; 3 seconds: -0.88 ± 3.48 vs. 3.38 ± 4.01, p=8.53e^-28^ ). Although the total response magnitude declined as the integration period shortened due to the reduced time for GCaMP6s signals to accumulate, the relationship between the AOSLO and Maxwellian view responses remained statistically significant. These results confirm that the lack of OFF cells identified in our earlier experiments arose because the AOSLO decrement stimulus was ineffective in driving strong positive responses in OFF cells.

Two general trends emerged from both the ON and OFF cell responses (**Figures 5A-B**). First, the responses to Maxwellian view stimulus were larger than the responses to the AOSLO stimulus. To quantify this, we calculated the amplitude (trough-to-peak distance) for each cell. The response amplitudes were significantly larger for Maxwellian view stimuli compared to AOSLO stimuli (1.11 ± 0.75 vs. 0.61 ± 0.29; p = 1.22e^-19^ ; **Figure 5E**). This suggests that ON and OFF cells utilized their full dynamic range more effectively in response to the Maxwellian view stimuli. Second, the Maxwellian view stimulus elicited more rectified responses than the AOSLO stimulus (i.e., ΔF/F increased but rarely decreased). We calculated a rectification index (RI) by dividing the peak value by the amplitude. An RI of 0.5 indicates equally weighted peak and trough values, while an RI of 1 indicates complete rectification with the range equal to the peak. The Maxwellian view stimulus responses were significantly more rectified than the AOSLO responses (0.90 ± 0.12 vs. 0.41 ± 0.26, n = 214; p = 4.47e^-87^ ). This increase in response rectification was largely attributable to the lack of negative decrement responses from ON cells (**Figures 5C-D**).

Taken together, these results confirm that AOSLO stimulus delivery is responsible for both the ON cells’ large negative decrement responses and the OFF cells’ low responsivity to decrement stimuli. The Maxwellian view stimulus elicited larger response amplitudes than the AOSLO stimulus, indicating that both ON and OFF cells were utilizing their full dynamic range more effectively for the Maxwellian view stimulus than for the AOSLO stimulus.

### 3.5. Hypothesized origin of unexpected responses to AOSLO visual stimuli

We next asked which aspect of AOSLO-mediated stimulation leads to the ON cells’ elevated baseline activity and large negative responses to decrements. A key consequence of raster-scanning is that the background around which contrast increments and decrements are presented is only uniform when integrated over a full cycle of the slow scanner, which sets the frame rate (Poonja et al., 2005). From the perspective of a single foveal cone —approximately 1 pixel in these experiments – the “background” light would not be a constant luminance but a periodic series of brief pulses of light. This temporal pattern could be a strong stimulus for ON cells, driving their continuous responses to the background light.

To test whether raster-scanned, AO-corrected light causes elevated baseline activity in ON cells, we asked whether the ON cells’ large negative responses to decrements (e.g., **Figure 2A**) could be replicated by simulating raster-scanning with the Maxwellian view. For the AOSLO used in these experiments, a full scanner cycle lasted ∼39.5 ms and the average dwell time over a single pixel was ∼35-60 ns. Thus, there are two key features to the temporal pattern of stimulation created by raster-scanning: the temporal frequency of 25.3 Hz and the duty cycle of 0.76-1.52e^-4^ %. We simulated this pattern by confining the light presented by the Maxwellian view during a 100% contrast decrement to a 2 ms pulse every 40 ms (**Figure 6A**). This created a periodic series of pulses with a temporal frequency of 25 Hz and a 2% duty cycle. Unfortunately, decreasing the duty cycle further to better approximate the brevity of AOSLO raster-scanning was prevented by the 500 Hz update rate of the LEDs in the Maxwellian view. Another difference was that the pulsed Maxwellian view stimulus was presented full-field while the AOSLO scanned a focused spot of light across the retina. However, the high speed of raster-scanning combined with the small size of foveal RGC receptive fields (most have single cone centers (Calkins et al., 1994) which were roughly 1 pixel at our FOV) suggest this difference may not be critical. Despite these differences, the pulsed Maxwellian view stimulus successfully replicated the ON cells’ large negative responses during the decrement stimulus. A comparison of the total stimulus response to the AOSLO decrement and the pulsed Maxwellian view decrements for 198 cells is shown in **Figure 6B**. ON cells that exhibited fluorescence decreases to the AOSLO decrement also showed fluorescence decreases to the pulsed Maxwellian view decrement.

**Figure 6.**
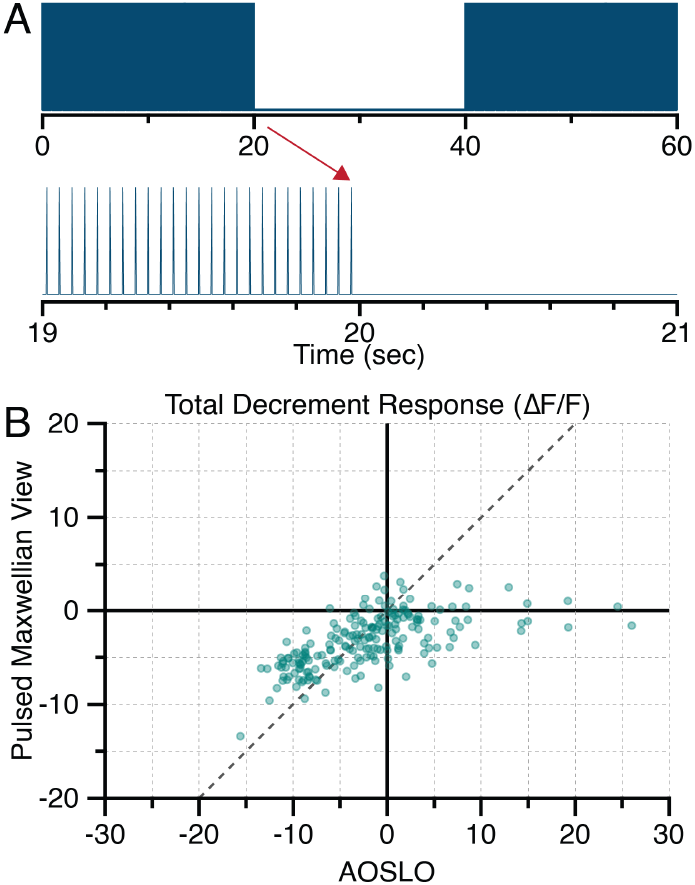
**(A)** Stimulus intensity for the pulsed Maxwellian view decrement stimulus in which the LEDs were on for just 2 ms every 40 ms (25 Hz with a 2% duty cycle). The stimulus in the bottom panel followed 2 minutes of adaptation to the pulsed “background” shown in the first 20 seconds. **(B)** Total stimulus responses for 198 cells to the AOSLO decrement stimulus and the “pulsed” Maxwellian view stimulus designed to imitate the temporal pattern of excitation created by raster-scanning (see text). The dark gray dashed line indicates where cells would be located if the two stimuli elicited identical responses.

Two unexpected features of the responses to the pulsed Maxwellian view decrements provided further insights. First, the pulsed Maxwellian view decrement elicited smaller negative responses in ON cells than the AOSLO decrement (bottom left quadrant of **Figure 6B**), indicating lower ON cell baseline activity levels in the pulsed Maxwellian view condition. Second, small negative responses were observed in OFF cells that had positive responses to the AOSLO decrement (bottom right quadrant of **Figure 6B**). This fluorescence decrease during the decrement implies the OFF cells had a non-zero baseline activity level prior to the decrement and were less rectified. This suggests that the asymmetric impact of AOSLO raster scanning on the baseline activity of ON vs. OFF cells was less extreme for the pulsed Maxwellian view stimulus. Specifically, OFF cells appeared to be less rectified because some baseline activity was required for negative responses during the decrement stimulus and ON cells appeared to have slightly lower, but still elevated, baseline activity. We propose this result arises from the increased duty cycle of the pulsed Maxwellian view stimulus.

The reduction in ON/OFF cell response asymmetry with an increased duty cycle may help bridge the gap between our findings and prior studies of the temporal frequency tuning of foveal RGCs using stimuli with a 50% duty cycle. Both ON and OFF foveal midget RGCs respond to 25 Hz sinusoidal temporal modulations with equal light:dark ratios (Solomon et al., 2002), which would predict similarly elevated baseline activity in both ON and OFF cells whereas raster-scanning at 25.3 Hz led to increased rectification in OFF cells. Future experiments would be needed to explore this idea further, but these different findings could be reconciled by the hypothesis that ON/OFF asymmetries become more pronounced as the light:dark ratio becomes increasingly asymmetric.

### 3.6. Adaptation to the onset of a photopic background light

The experiments thus far have drawn conclusions about elevated baseline activity from stimulus-response functions and the presence of large negative responses in normally-rectified RGCs. To directly test the hypothesis that the raster-scanned background light alters the baseline activity in ON and OFF cells, we investigated the time course of adaptation to the onset of the background light used in **Figures 2, 3** and **5**. We refer to this as a “step stimulus” as it reflects a step increase in stimulus intensity from a low photopic light level to a high photopic light level. Importantly, a dim adapting light was present even in the pre-stimulus time due to inevitable leak light from the 561 nm laser’s AOM and the presence of the 796 nm reflectance imaging light (see Methods).

If raster-scanning AO-corrected light drives high baseline activity in ON cells, then the AOSLO step stimulus should produce stronger, more sustained responses in ON cells than the Maxwellian view step stimulus. A related prediction is that ON cells will adapt to the step stimulus presented by the Maxwellian view, but not the AOSLO. To test these predictions quantitatively, we computed the absolute “peak response” during the step stimulus and the “final response”, which was defined as the median ΔF/F during the final 5 seconds of the step. The peak response established the magnitude of change in ΔF/F during the step stimulus while the final:peak response ratio quantified the extent of response adaptation (see Methods). Based these metrics, the ON cells’ peak response is predicted to be higher for the AOSLO step stimulus, and the final response is predicted to be a greater percentage of the peak response (i.e., a “final:peak response ratio” near 1).

Comparison of the population responses to the AOSLO and Maxwellian view step stimuli revealed substantial differences in response magnitude and duration (n = 298 cells; **Figure 8A**). **Figure 8B** compares the peak response and the final:peak response ratio for the same 298 cells. Representative responses to the AOSLO and Maxwellian view step stimuli for four ON and four OFF cells are shown in **Figure 8C-D**, with the final:peak ratios are listed for reference.

**Figure 7.**
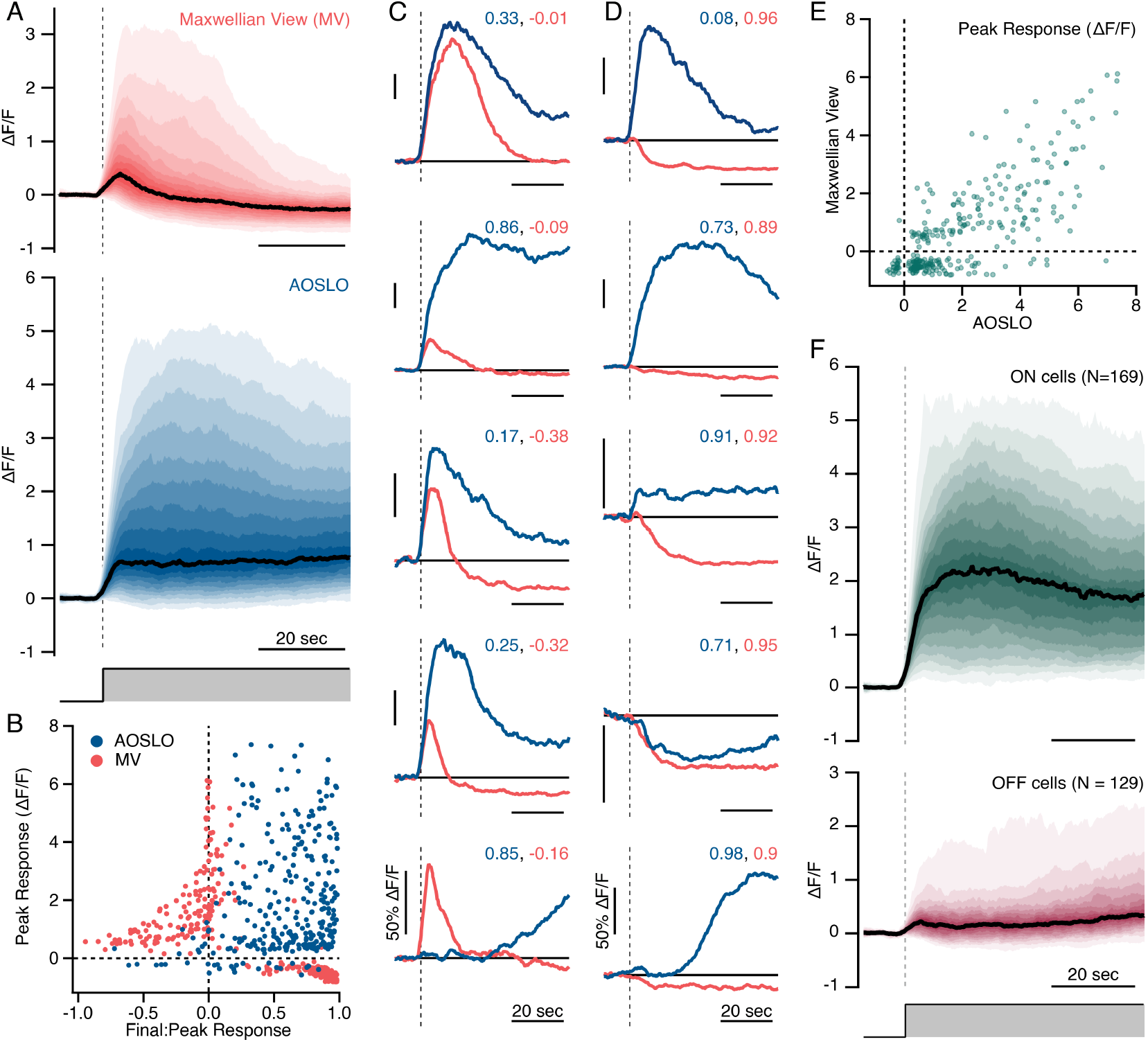
**(A)** Fancharts showing the population-level response of 298 cells to step stimuli presented by the LEDs (red) and the AOSLO (blue). The black lines show the median response and the shaded regions show the 5-95th percentiles in increments of 5. Dashed lines indicate the time of step onset. **(B)** The peak response and final:peak response ratio for 298 cells. Each cell is plotted twice, once in red showing the response to the Maxwellian step stimulus and again in blue showing the response to the AOSLO step stimulus. **(C-D)** Representative responses from four ON cells (C) and four OFF cells (D). To provide intuition for the interpretation of the final:peak responses in B, the final:peak values are included for each representative trace. As in the previous figures, red traces and text are used for the Maxwellian view step stimulus responses, blue traces and text are used for the AOSLO step stimulus, and gray vertical dashed lines indicate the time of step onset. All horizontal scale bars are 20 seconds and all vertical scale bars are 50% ΔF/F. **(E)** Direct comparison of the peak responses to AOSLO and Maxwellian view step stimuli for 298 cells. **(F)** Fan charts showing the population-level responses of 169 ON cells and 129 OFF cells to the AOSLO step stimulus. The black lines show the median response and the shaded regions show the 5-95th percentiles in increments of 5. Dashed lines indicate the time of step onset.

**Figure 8.**
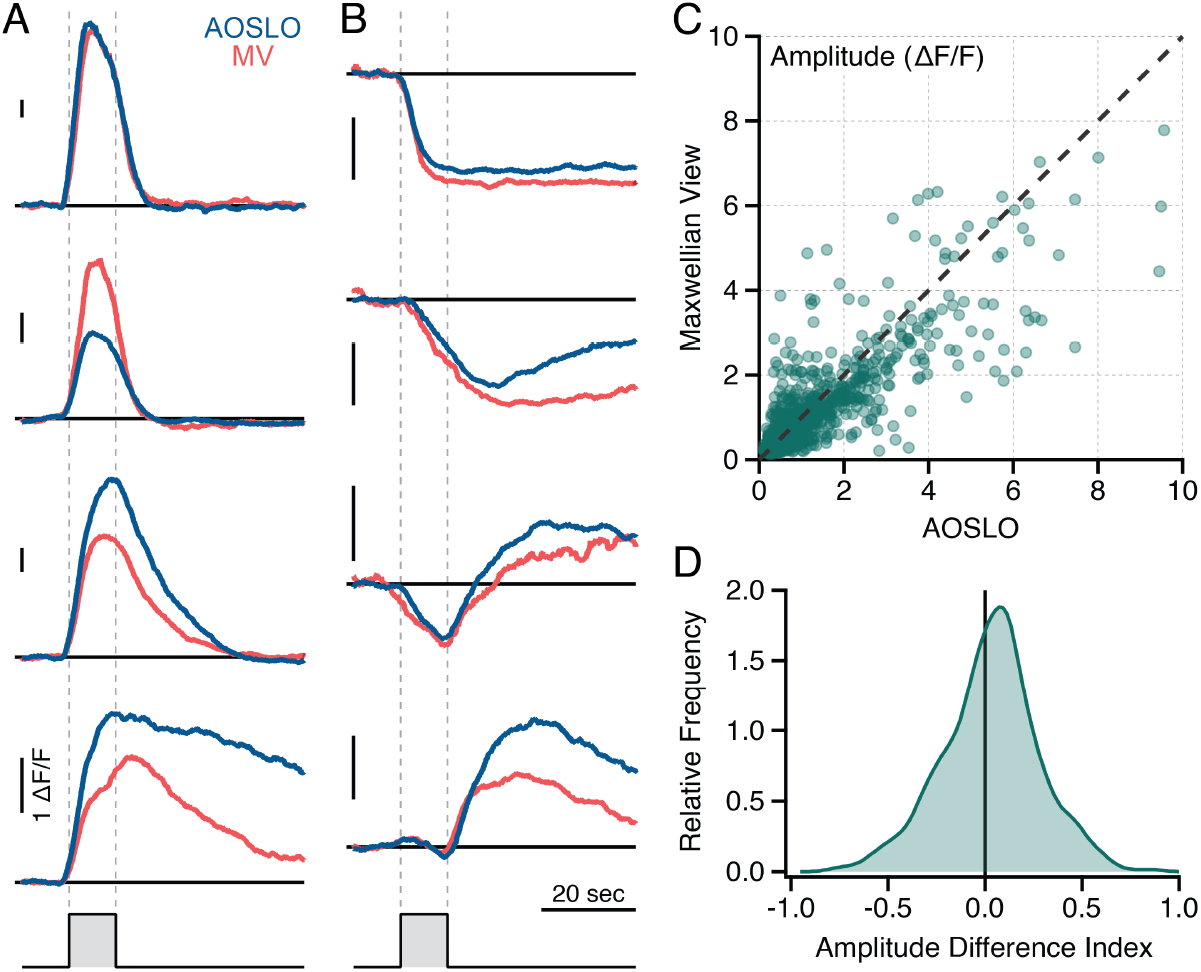
**(A)** Representative responses from four ON cells to 10-second intensity increments presented through the AOSLO (blue) and with the Maxwellian view (red). The vertical dashed lines mark the onset and offset of the intensity increment. All horizontal scale bars are 20 seconds and all vertical scale bars are 100% ΔF/F. **(B)** As in A, but showing representative responses from 4 OFF cells. **(C)** Comparison of the trough-to-peak response amplitudes for responses to the AOSLO and Maxwellian view intensity increments for 882 cells. **(D)** Kernel density estimate of the distribution of amplitude difference indices (ADIs; see text).

Responses to the Maxwellian view step stimulus were largely consistent with light adaptation following an increase in mean light level (Kuffler et al., 1957; Barlow and Levick, 1969; Yeh et al., 1996), albeit with a prolonged time course due to the slow decay of GCaMP6s. Most ON cells with positive peak responses exhibited final:peak ratios between 0.1 and -1 (**Figure 8B**), indicating that after a transient increase in fluorescence, the response returned near or below the pre-stimulus baseline fluorescence. Most OFF cells exhibited a monotonic decrease in fluorescence that stabilized over 20–40 seconds. In this way, ON and OFF cells could be classified by the sign of their peak response to the Maxwellian view step stimulus (**Figure 8B**). The bimodal distribution of peak responses was statistically significant (Hartigan’s dip test, p = 0.001), and a two-component Mixture of Gaussians model identified two groups (1.677 ± 1.457, n = 158 and -0.508 ± 0.154, n = 140).

In contrast, responses to the AOSLO step stimulus were larger and more sustained than those to the Maxwellian view step stimulus. Notably, very few cells exhibited negative peak responses to the AOSLO step stimulus, as most cells were located to the right of the vertical dashed line in **Figure 8B**. This pattern is further highlighted in **Figure 8E**, which directly compares the peak responses to the AOSLO and Maxwellian view step stimuli across all 298 cells. Interestingly, most OFF cells that showed a fluorescence decrease in response to the Maxwellian view step stimulus instead exhibited an increase in fluorescence to the AOSLO step stimulus. Overall, the AOSLO step responses were inconsistent with traditional light adaptation, suggesting that the retina does not process the AOSLO step stimulus as a change in mean light level.

We next asked whether ON and OFF cells responded differently to the AOSLO step stimulus. We used the Mixture of Gaussians model above to classify ON and OFF cells based on their peak responses to the Maxwellian view step stimulus. Visual inspection of the individual responses revealed 12 misclassified ON cells with late decreases in fluorescence that were slightly larger than the initial transient increase. These cells were reassigned to the ON group. **Figure 8F** shows the median responses and percentiles for ON and OFF cells to the AOSLO step stimulus. The majority of both ON and OFF cells showed some increase in fluorescence during the AOSLO step stimulus (**Figure 8E**); however, the ON cells’ responses were much larger and more sustained. Two-sample Kolmogorov-Smirnov tests confirmed that these trends were significant: ON cells exhibited larger peak responses (2.98 ± 1.88, n = 169 vs. 0.79 ± 0.16, n = 129; p = 3.60e^-24^ ) and final responses (1.92 ± 1.47, n = 169 vs. 0.54 ± 0.83, n = 129; p = 8.23e^-18^ ) compared to OFF cells. The slight increase inΔF/F near the end of the stimulus in the median OFF cell response was driven by a population of rarer cells with large, late increases in ΔF/F (e.g., the bottom trace of **Figure 8D**).

These results confirm that raster-scanned background lights induce elevated activity in the ON pathway, but not in the OFF pathway. Given the temporal pattern of excitation created by raster scanning, step stimuli presented through the AOSLO primarily represent changes in temporal contrast. From the perspective of an individual cone photoreceptor, even the 0% contrast control stimulus that was intended to provide a constant background light actually consists of a series of brief, high-intensity pulses, making it a stimulus with high temporal contrast.

### 3.7. Responses to increments when the background intensity is minimized

If the raster-scanned background light is a key factor in the unexpected ON and OFF cell responses to increment and decrement stimuli, then the AOSLO and Maxwellian view stimulus responses will be more comparable when the background intensity is minimized. We tested this prediction by comparing the responses to a 10-second intensity increment presented without a mean background. Note that while these conditions minimize the impact of raster-scanning, a low photopic background remains present due to the reflectance imaging light and inevitable leak of the 561 nm laser light through the AOM (Domdei et al., 2018). These conditions remain relevant to the many AOSLO psychophysics experiments performed with similar background light levels (Tuten et al., 2012; Harmening et al., 2014; Sabesan et al., 2016).

**Figures 8A-B** show representative ON and OFF cell responses to the intensity increments presented by both stimulators. Although no clear trends were apparent in the representative traces, quantification of the response amplitudes revealed slightly larger responses to the AOSLO stimulus (1.34 ± 1.39 vs. 1.19 ± 1.13, n = 882; p = 0.016; **Figure 8C**). To further assess the per-cell differences, we calculated an amplitude difference index (ADI), defined as the difference between the AOSLO and Maxwellian view response amplitudes divided by their sum. The ADI ranges from -1 to 1, where positive values indicate a larger AOSLO response, and the magnitude reflects the extent of the amplitude difference (**Figure 8D**). Across 882 cells, the ADI was 0.29 ± 0.249 with a median value of 0.041, indicating that the AOSLO response amplitudes were generally larger than those for the Maxwellian view. Despite minimizing the background light, this result could still stem from the temporal pattern of excitation created by the AOSLO’s raster-scanning. Unlike the Maxwellian view increment, which maintains a remained constant intensity, the AOSLO increment consists of a series of brief, high intensity pulses that may drive more sustained responses and less adaptation over the 10 second stimulus duration.

Despite the small, but significant, difference in response amplitude, the AOSLO and Maxwellian view responses appeared qualitatively similar (**Figure 9A**). To quantify the similarity between the response shapes, we computed the correlation coefficients between the normalized responses to the AOSLO vs. Maxwellian view intensity increments for 882 ROIs from three retinal regions in two macaques. The correlation was strong, with an average correlation coefficient of 0.614 ± 0.436 and a median correlation coefficient of 0.804. An empirical cumulative distribution function of the correlation coefficients is shown in **Figure 9A**. For reference, we repeated this analysis with the decrement-increment stimulus (0.08 ± 0.51, n = 220) and the control stimulus (median: 0.023, mean ± SD: 0.011 ± 0.320, n = 690). Both were weakly correlated, although the spread of correlation coefficients was much larger for the decrement-increment stimulus, consistent with the ON cells’ opposite responses to AOSLO vs. Maxwellian view decrements seen in **Figure 5**.

**Figure 9.**
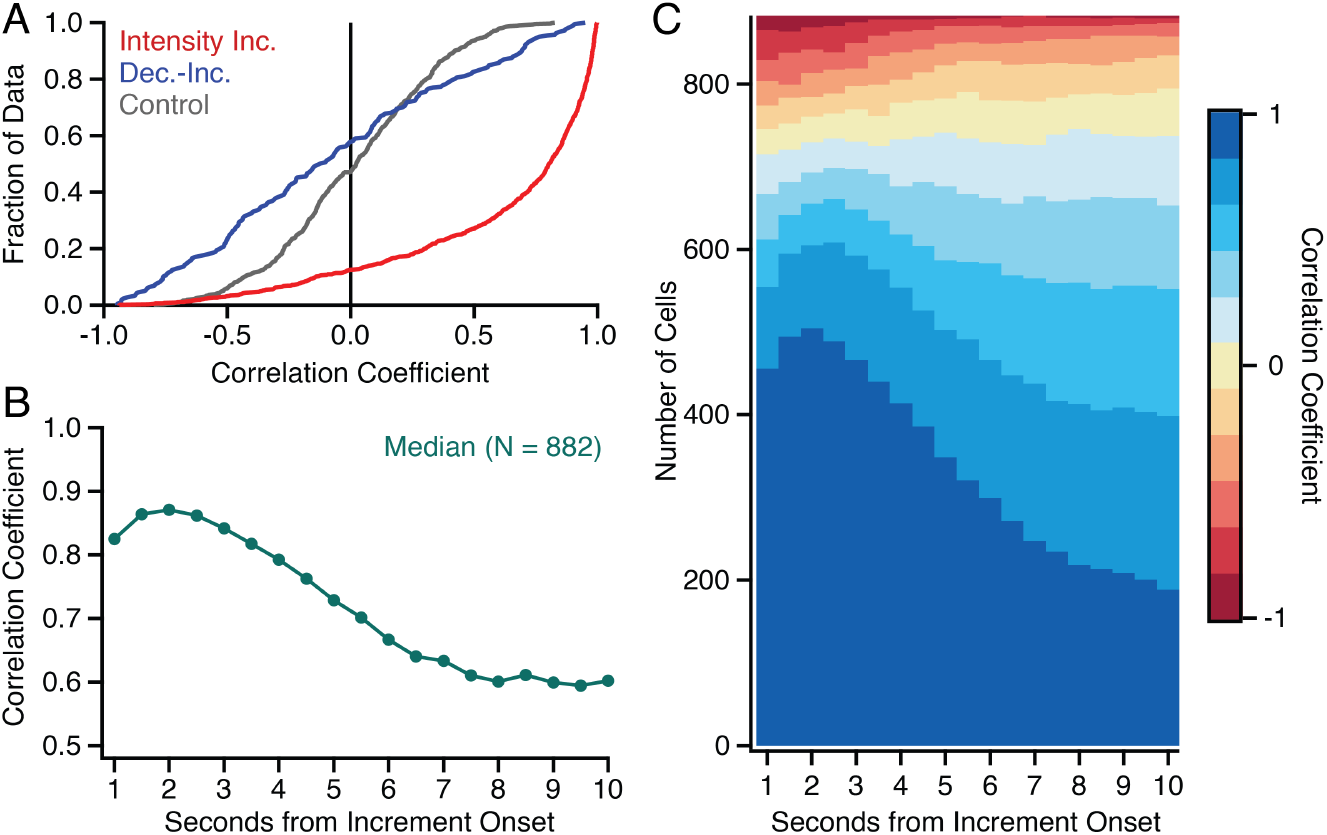
**(A)** Empirical cumulative histograms of the correlation coefficients between normalized responses to AOSLO and Maxwellian view stimulus delivery for three different stimuli: intensity increments (red; **Figure 8**), the decrement-increment stimulus (blue; **Figure 5**) and a control stimulus at a photopic background light level. The full stimulus time course was used, beginning at stimulus onset. **(B)** The median correlation coefficient for AOSLO and Maxwellian view intensity increments calculated over various time periods, from 1-10 seconds after stimulus onset. **(C)** The correlation coefficients obtained at each time period for 882 cells, sorted from bottom to top in descending order.

Taken together, these data indicate that calcium responses to the AOSLO and Maxwellian view intensity increments were qualitatively comparable. It is worth noting that many differences were likely exaggerated by the long 10-second increment. The intensity increment correlation coefficients in **Figure 9A** were based on the full response (the 10-second increment and 45 seconds following the increment). By contrast, the increments presented in AOSLO-mediated psychophysics are generally <1 second long (e.g., 500 ms (Sabesan et al., 2016)). To obtain results more comparable with these stimuli, we next analyzed the responses. **Figure 9B-C** show the median and population-level correlation coefficients over different time periods following the onset of the intensity increment. For example, the correlation coefficient at 1 second was calculated based on the responses during the first second following stimulus onset. The median correlation coefficient was highest 1.5-2.5 seconds after stimulus onset. Taken together, these results indicate that the brief intensity increments used in AOSLO-mediated psychophysics are performed under conditions that minimize the influence of raster-scanning on retinal physiology.

## 4. Discussion

Here we assessed the responses of the retinal ON and OFF pathways to stimuli presented through an AOSLO and via a 3-LED Maxwellian view stimulator. Despite numerous technical differences between *in vivo* calcium imaging and *ex vivo* electrophysiology, we found qualitative agreement between the responses to the Maxwellian view stimuli and previously published work. However, unexpected responses were observed for some stimuli delivered through the AOSLO, suggesting that the temporal pattern of excitation created by raster-scanning AO-focused visible light alters the response properties of ON and OFF RGCs. Specifically, a consequence of raster-scanning is that the photopic “background light” is not a spatiotemporally constant illumination, but rather a strong periodic stimulus with high temporal contrast. Our results are consistent with the hypothesis that altered baseline activity due to the photopic raster-scanned background lights lead to high baseline activity in ON cells and increased rectification in OFF cells. This increased depolarization of ON cells and hyperpolarization of OFF cells can explain the unexpected contrast increment and decrement responses produced by photopic contrast modulations delivered through the AOSLO.

### 4.1. Impact of raster-scanning differs for ON and OFF cells

A working model of the circuit mechanisms underlying ON and OFF RGC responses to photopic raster-scanned stimuli is essential for an informed interpretation of how our results impact AOSLO-mediated psychophysics and physiology. The mechanism driving ON cell responses appears relatively straightforward: raster-scanning elevates baseline activity, shifting their dynamic range toward decrements and limiting their capacity to signal increments. This conclusion is supported by the ON cells’ sustained response to the AOSLO step stimulus (**Figure 7**) and large negative responses to high contrast decrements (**Figures 2-3**), neither of which occurred for comparable stimuli presented with the Maxwellian view (**Figure 5**). Additional evidence comes from the ON cells’ shallow increment CRFs (**Figures 2A**) and declining increment responses as background light levels increased (**Figure 4**); both are consistent with the ON cells responding near saturation to raster-scanned backgrounds.

In OFF cells, greater rectification was observed, which minimized responsivity to all but the highest contrast decrements (**Figure 2B**). As background light level increased, reduced responsivity was observed even for 100% contrast decrements (**Figure 4C**). Furthermore, our data underestimates the extent to which OFF pathway sensitivity is reduced. When OFF RGCs were identified by their responses to AOSLO decrements, ON cells far outnumbered OFF cells (e.g., 3.1:1 in **Figure 3B**; see also the asymmetry in **Figures 2B** and **4B**). This discrepancy was far smaller when ON and OFF cells were classified based on their responses to the Maxwellian view stimuli (**Figure 7F**).

Why do ON cells exhibit larger, more sustained responses to the raster-scanned backgrounds than OFF cells? One possibility is that ON cells simply respond more robustly to the brief intensity pulses generated by raster-scanning. However, robust responses to brief intensity flashes are observed in OFF cells, albeit following a period of inhibition corresponding to the ON cells’ initial response (Kuffler, 1953; Büttner et al., 1975). This alternation between ON and OFF responses points to a second complementary possibility: increased activity in the ON pathway may inhibit the OFF pathway. This explanation is compelling because ON→OFF pathway inhibition is key in establishing the asymmetric contrast responses and differing baseline activities of ON and OFF RGCs (**Figure 3A**) (Zaghloul et al., 2003; Rentería et al., 2006; Margolis and Detwiler, 2007; Molnar and Werblin, 2007; Liang and Freed, 2010). The underlying circuit is depicted in **Figure 10**.

**Figure 10.**
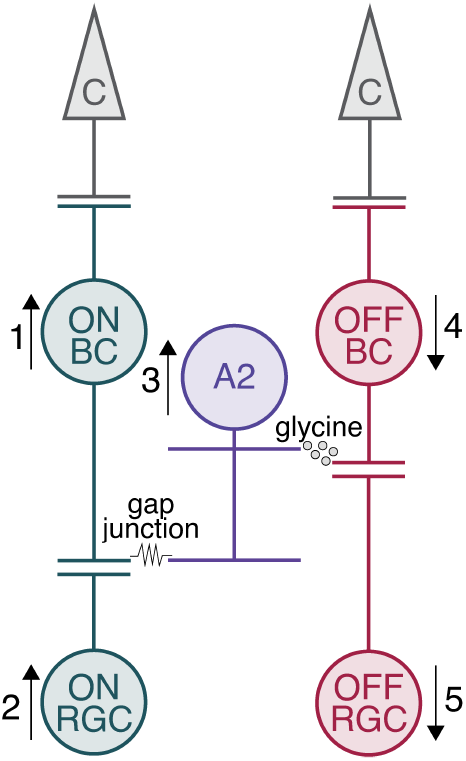
Retinal mechanisms underlying ON-OFF asymmetry in maintained discharge under photopic light levels provide a working model for the impact of raster-scanned background light. At rest, ON cells have some baseline discharge (e.g., 12 spikes/sec; (Percival et al., 2022)) while OFF RGCs fall near 0 spikes/sec. This asymmetry arises because the ON bipolar cells are slightly depolarized (1) at rest and releasing glutamate to the ON RGCs (2). The depolarization of ON bipolar cells spreads through gap junctions to the AII amacrine cells. When depolarized, the AII amacrine cells increase glycine release onto the OFF bipolar cells (3). Glycine, an inhibitory neurotransmitter, hyperpolarizes the OFF bipolar cells, reducing their glutamate release onto OFF RGCs (4-5) at the same time that the depolarized ON bipolar cell increases glutamate release onto ON RGCs (1-2). Any stimulus that further drives the ON pathway – such the brief pulses of light created by raster-scanned background lights – will further depolarize the ON bipolar cells and ON RGCs (1-2) and increase the ON*→*OFF pathway inhibition (3) through this circuit, resulting in greater rectification of the OFF pathway neurons (4-5).

At rest, ON RGCs exhibit higher maintained activity which reflects a baseline level of depolarization in presynaptic ON bipolar cells. This depolarization spreads from ON bipolar cells to AII amacrine cells via gap junctions. The resulting depolarization of AII amacrine cells increases their release of glycine, an inhibitory neurotransmitter, onto post-synaptic OFF bipolar cells, including the OFF midget bipolar cells presynaptic to the most common foveal OFF RGC in primates (McLaughlin et al., 2021; Percival et al., 2022). The resulting hyperpolarization of OFF bipolar cells rectifies their glutamate release onto post-synaptic OFF RGCs, leading to low baseline firing rates and decreased responsivity to low contrast decrements. This circuit establishes the OFF pathways at scotopic light levels and helps maintain the ON and OFF pathways’ asymmetric contrast sensitivities at photopic light levels (Demb and Singer, 2012). The latter role may be one reason why AII amacrine cells are present in the primate fovea, even where rods are absent (Strettoi et al., 2018).

Any stimulus that drives the ON pathway, like a contrast increment or the brief pulses of light created by AOSLO raster-scanning (**Figure 6A**), increases ON bipolar cell and ON RGC activity and could also be expected to hyperpolarize OFF bipolar cells and further rectify the responses of OFF RGCs. This explanation could also account for the increased rectification of decrements under 100% contrast seen in the OFF cell CRFs in **Figure 3B**. Even a 75% contrast decrement still contains brief, periodic flashes of light capable of driving the ON pathway and hyperpolarizing the OFF pathway.

While voltage clamp measurements of inhibitory and excitatory currents along with pharmacology would provide more direct support for AII amacrine cell’s involvement, the circuit in **Figure 10** is a strong candidate for mediating the observed ON-OFF asymmetries to raster-scanning. First, it does not require novel circuitry, but instead relies on an exaggerated effect of the well-established pathway for establishing the OFF pathway’s rectification (Zaghloul et al., 2003; Margolis and Detwiler, 2007; Liang and Freed, 2010; Percival et al., 2022). Second, feedback inhibition to OFF bipolar cells is not incompatible with the minimal direct, feedforward inhibition reported in foveal midget RGCs (Sinha et al., 2017), which rules out many alternative inhibitory circuits. Third, this circuit has been implicated in similar ON-OFF asymmetries that arise in response to high-frequency electrical stimulation by retinal prosthetics. Much like the high-frequency pulses created by raster-scanning, electrical stimulation is more effective for ON cells than OFF cells (Im and Fried, 2016; Lee and Im, 2019). An increase in glycinergic inhibition to OFF bipolar cells through the circuit shown in **Figure 10** is thought to contribute to the reduced excitability of OFF RGCs to electrical stimulation (Carleton and Oesch, 2024).

### 4.2. Impact on AOSLO-mediated psychophysics and physiology

How do these findings affect the interpretation of AOSLO-mediated psychophysics? The most common stimulus paradigms involve brief intensity increments targeting small retinal regions or even single cone photoreceptors (Tuten et al., 2012; Harmening et al., 2014; Sabesan et al., 2016; Vanston et al., 2023). The short stimulus duration and lack of a strong raster-scanned background both likely minimize the impact of raster-scanning considerably (**Figures 8-9**). Additionally, increased awareness of light leakage from modern AOMs have led to new strategies for further suppressing unintentional raster-scanned background lights, which could impact both RGC responses and visual perception. For example, Domdei and colleagues recently demonstrated high contrast AOSLO visual stimuli can be obtained by chaining two AOMs to decrease leaked light (Domdei et al., 2018). Many psychophysics experiments also present adapting lights with a Maxwellian view to further reduce the effects of raster scanned background light (Tuten et al., 2017; Schmidt et al., 2018a; Greene et al., 2024).

Raster-scanning would be expected to play a larger role in experiments presenting increments and decrements relative to high photopic backgrounds. Our data predict that, under these conditions, decrements to be more salient and exhibit lower detection thresholds than increments. Indeed, many AOSLO psychophysics experiments create spatial stimuli by switching off the scanning laser, thereby creating contrast decrements (Ratnam et al., 2017). In these experiments, the goal is generally to measure visual acuity or chromatic aberration and achieving a specific balance of ON vs. OFF pathway activation may not be essential. However, when considering the potential retinal mechanisms contributing to these decrement stimuli, it will be important to keep in mind that both ON and OFF cells signaling each decrement (**Figure 2**). Moreover, depending on the decrement’s contrast and the background light level, ON cells may be contributing just as much, if not more, than OFF cells (**Figures 3** and **4**).

It is interesting to note that, although the circumstances created by raster-scanning are uncommon, their impact largely exaggerates a pre-existing bias towards decrements (Krauskopf, 1980; Yeh et al., 2009; Cooper and Norcia, 2015). This bias begins in the cone photoreceptors (Angueyra et al., 2022) and is reinforced by the asymmetric contrast sensitivities of the retinal ON and OFF pathways (Chichilnisky and Kalmar, 2002; Zaghloul et al., 2003). As can be seen in **Figure 3A**, both ON and OFF RGCs respond to decrements, while only ON RGCs respond to increments, potentially enhancing decrement detection. At the low contrasts that dominate natural scenes (Tadmor and Tolhurst, 2000), ON cells are thought to play a larger role in signaling decrements than OFF cells.

### 4.3. Mitigating the impact of raster-scanning for AOSLO-mediated visual stimuli

There are two raster-scanning hardware control parameters that could alter the impact of raster-scanning on RGC physiology: the temporal frequency and the duty cycle. The extent to which these two parameters can be changed is currently limited by the hardware used for raster-scanning. Reducing the field of view in order to increase the dwell time of the scanned light over each photoreceptor would yielding slight increases in the duty cycle. However, our pulsed Maxwellian view experiment in **Figure 6** indicated that small increases in duty cycle could reduce the imbalance between ON and OFF cells to some degree by increasing the baseline firing rate in OFF cells and reducing the baseline firing rate in ON cells. In other words, changing the duty cycle to achieve a more uniform light:dark ratio may reduce the imbalance between the ON and OFF pathways, but will be unlikely to produce conditions comparable to more traditional visual stimulators unless the temporal frequency is also increased.

Indeed, increasing the temporal frequency of raster-scanning may be necessary to mitigate its impact on RGCs physiology. In calcium imaging experiments with *ex vivo* retinal tissue, visual stimuli are often presented during the back-scan of the excitation laser; for example, at 500 Hz with a 25% duty cycle (Euler et al., 2009; Franke et al., 2019). At such a high frequency, the impact of the duty cycle is likely negligible. In choosing an appropriate temporal frequency, it will be important to consider frequencies much higher than the standard temporal resolution limits measured psychophysically. First, many RGCs and their downstream projections do respond to higher temporal frequencies than can be resolved perceptually, with some retinorecipient neurons in the lateral geniculate nucleus responding up to 100 Hz (Tailby et al., 2007). Second, temporal resolution limits are much higher when periodic stimuli have extreme light:dark ratios (Cobb, 1934; McNemar, 1951), similar to that created by raster-scanning AO-focused light.

Most AOSLOs achieve raster-scanning with a 14-16 KHz horizontal resonant scan mirror and a 25-30 Hz vertical galvanometric scan mirror (Poonja et al., 2005). In these systems, some capacity exists to achieve higher temporal frequencies without system modifications. For example, FOV width can be traded for increased temporal frequency; by halving the period of the slow galvanometer scanner, a single frame could be completed in half the time. With the most commonly-used resonant and galvanometric scanners, this approach can be used for raster scanning at 100-200 Hz. However, whether the necessary reduction in FOV width is feasible will be highly experiment-dependent. For example, eye tracking and stimulus stabilization may be compromised, particularly in participants with large natural eye movements, and the spatially-offset stimulus/imaging configuration in **Figure 1** would be challenging.

### 4.4. Alternative stimulation strategies for AOSLO

AOSLO enables a unique combination of highresolution retinal imaging, eye tracking, and stimulus delivery. In many cases, AOSLO has enabled measuring structure-function relationships at a microscopic scale that is not possible with other techniques (Sabesan et al., 2016). For example, the unique insights AOSLO has provided into the pathology of rare, understudied inherited retinal degenerations likely outweigh any concerns about the impact of raster-scanning (Duncan and Carroll, 2022). However, the key advantages provided by stimulation through the AOSLO (AO-corrected spatial resolution, precise stabilization and localization of stimuli on the cone mosaic) must be weighed against the impact of raster-scanning AO-corrected visual light on RGC responses and other limitations (low frame rates and small FOVs).

One straightforward technique demonstrated here and in our prior work (Godat et al., 2022, 2024) is the presentation of spatially-uniform stimuli using a Maxwellian view stimulator. In our experiments with anesthetized and paralyzed macaques, some residual eye motion remains due to head motion during respiration; however, lack of stabilization was not an issue. By using a larger stimulus area than necessary, receptive fields remained in a region of spatially-homogeneous illumination (see Methods). A similar approach has been utilized to provide chromatic adapting backgrounds during psychophysics experiments using AOSLO to deliver small spots of light to individual cones (Tuten et al., 2017; Schmidt et al., 2018b). Where spatial stimuli are not necessary, this is a straightforward and effective solution with few drawbacks and superior spectral and temporal control.

Maxwellian view stimulators have also been used to present spatial stimuli in AO-mediated psychophysics experiments (Wang et al., 2023; Greene et al., 2024). While this strategy has yet to be used with retinal physiology, it represents a valuable potential future direction. An important caveat is that most AOSLO retinal imaging is performed with a dilated pupil which will compromise the spatial resolution of stimuli (Fitzpatrick et al., 2024; Cobb, 1915; Campbell and Gregory, 1960). A recently reported monitor-based stimulator for AOSLO retinal imaging overcame this obstacle by placing an artificial 3 mm pupil in the stimulator arm (Moon et al., 2024). While these stimuli do not benefit from the precision provided by AO, neither do the visual stimuli we encounter in natural vision – in this sense, stimuli that are not AO-focused are more naturalistic and, for many research questions, further resolution may not be necessary. Experiments that do require AO-corrected stimuli may benefit from combining AOSLO with an AO visual stimulator or an AO flood-illuminated ophthalmoscope (Marcos et al., 2022; Teverovsky et al., 2024).

### 4.5. Impact on functional imaging with AOSLO stimulus delivery

While these results present a challenge for calcium imaging experiments aiming to collect data comparable to more traditional *ex vivo* electrophysiology techniques, it is less clear whether they diminish the insights gained from experiments using AOSLO-mediated stimulus delivery. The vast majority of psychophysics and retinal physiology is performed with contrast modulations around a spatiotemporally uniform background (e.g., drifting gratings and other sinusoidal contrast modulations on a gray background). These conditions minimize the number of variables affecting the neuronal response properties and keeps adaptation state fully under experimenter control. Yet constant adapting backgrounds are quite unlike natural vision, where the eye is constantly in motion and encountering intensity changes differing over several orders of magnitude (Frazor and Geisler, 2006; Rieke and Rudd, 2009). Regardless, the lack of control over the temporal characteristics of AOSLO raster-scanning and the reduced response amplitudes observed in **Figure 5E** both raise concerns for many types of retinal physiology experiments.

While several of the spatial stimulators discussed above have provided valuable solutions for human AO psychophysics experiments, several technical obstacles remain for their application to functional imaging in anesthetized animal models. For example, both approaches rely on the participant feedback to ensure the stimulus is in-focus at the beginning of an experiment; this feedback is not easily achieved with non-human physiology given that anesthesia and paralysis are currently both required to optimize the SNR of calcium responses. The lack of stabilization may also be problematic, if the goal is to mirror the conditions under which *ex vivo* retinal physiology data is collected. The vast majority of standard retinal physiology stimuli and analyses assume a stationary retina with stabilized stimuli; where eye movements are incorporated, their trajectory and statistics are controlled by the experimenter. However, AOSLO psychophysics and other techniques have underscored the importance of natural eye movements in visual processing and the recent demonstration of precise receptive field mapping in the presence of eye motion suggests creative new analyses may be able to overcome a lack of stabilization (Yates et al., 2023). Alternatively, recently developed techniques for high-precision eye tracking based on imaging the pupil instead of the cone mosaic may soon provide the necessary feedback for stimulus stabilization (Kowalski et al., 2021; Wu et al., 2023).

### 4.6. Limitations

Each retinal physiology technique has a unique set of strengths and weaknesses that influence stimulus choice. As a result, direct comparisons between the results obtained by each technique are never perfect. The electrophysiology data we compared our results with was largely obtained with full-field spatial noise (multielectrode recordings (Chichilnisky and Kalmar, 2002)), brief contrast flashes and uniform temporal noise typically restricted to the center receptive field (singlecell retinal physiology (Zaghloul et al., 2003), or drifting gratings with spatial and temporal frequencies optimized for each receptive field (single-cell LGN physiology (Solomon et al., 2002)). For *in vivo* calcium imaging, we had to adjust our stimulus parameters and analyses to accommodate the specifics of our approach. For example, longer stimulus durations are common with *ex vivo* calcium imaging experiments (Baden et al., 2016; Franke et al., 2017) due to the slow kinetics of calcium indicators and low sampling rates of data acquisition. However, we extended stimulus duration much further in the present study to also accommodate the decreased SNR associated with imaging through the eye’s optics with low excitation light levels to prevent retinal damage. With this approach, we maximized detected responses but lacked the ability to detect spikes or calcium transients. The lack of single spike/transient resolution prevented more nuanced analyses and may have obscured weaker responses. As discussed above, one area where we suspect this may have limited our investigation is in the detection of OFF RGC responses to contrast decrements presented through the AOSLO. Strategies for addressing these limitations in the future include technical solutions for improving the SNR of *in vivo* retinal imaging and the use of optimized calcium or voltage indicators (Lu et al., 2023; Platisa et al., 2023; Zhang et al., 2023).

It is also worth noting that our experiments were performed at a relatively large FOV that exceeded the isoplanatic patch (Bedggood et al., 2008), which often serves as an upper bound on FOV in AO experiments. The impact of raster-scanning may vary with FOV size and with receptive field position within the FOV due to variations in the duty cycle, or light-dark ratio (amount of time a cone is exposed to light during each scanner cycle). For example, decreasing the FOV increases the number of pixels per cone and spreads excitation out over multiple scanned lines. As a result, the time when light is scanned across each cone increases and potentially approaches a uniform illumination. Similarly, the impact of raster-scanning could vary with receptive field position within the field of view, because the scanned light travels faster at the center of the FOV and slower at the edges due to the sinusoidal profile of the resonant scanner (Yang et al., 2015). While we did not observe an effect of receptive field position in our data, we cannot rule out a larger impact with smaller FOVs.

## 5. Conclusions

Despite considerable technical differences, *in vivo* calcium imaging can produce increment and decrement responses that are qualitatively comparable to *ex vivo* electrophysiology, but only when stimulus delivery was independent of the AOSLO. When stimuli were presented through the AOSLO, the temporal pattern of excitation created by raster-scanning AO-corrected light altered RGC response properties by altering their baseline activity (**Figures 6** and **7**). The magnitude of the impact on RGC physiology was stimulus-dependent and differed between ON and OFF RGCs. Our results led to a working model for the underlying circuit mechanisms mediating the responses of the most common ON and OFF RGC types. While the impacts of raster-scanning cannot be fully eliminated, they can be mitigated with stimulus choice. Our data indicate brief intensity increments minimize response discrepancies for raster-scanned, AO-corrected visual stimuli (**Figures 8** and **9**). Experiments using contrast modulations against a raster-scanned background light should ensure the mean light level is no brighter than necessary (**Figure 4**). Under these conditions, detection and discrimination of increments and low contrast decrements may be compromised (**Figure 3**).

## Acknowledgements

This work was supported by the National Eye Institute under F32-EY032318, P30-EY001319, R00-EY035323, R01-EY031467, R01-EY021155, and T32-EY007125, the Air Force Office of Scientific Research under FA-9550-22-1-0044 and FA-9550-22-1-0167, and Research to Prevent Blindness through an unrestricted grant to the Flaum Eye Institute. We thank Amber Walker and Jennifer LaPorta for animal anesthesia during imaging, the vector core at the Perelman School of Medicine of the University of Pennsylvania, the Genetically-Encoded Neuronal Indicator and Effector (GENIE) project, and the Janelia Research Campus of the Howard Hughes Medical Institute.

## Disclosures

QY has patents with the University of Rochester, Canon Inc., and the University of Montana (#9226656, #9406133, and #9454084) and has consulted for Oculus VR and Boston Micromachine Corporation. SSP has a patent application filed with the University of Washington (#17/612061).

## Data Availability Statement

The data and analysis code underlying the results presented in this manuscript will be made publicly available upon publication at https://github.com/sarastokes/aoslo-retinal-physiology.

